# Different tissues in the maternal-fetal interface harbor distinct microbiomes showing associations related to their anatomical position or function

**DOI:** 10.1101/2022.11.29.518443

**Authors:** Xiaopeng Li, Wei Jiang, Lijuan Dai, Guihong Liu, Bolan Yu, Min Fang

## Abstract

The human placenta was thought to be sterile in healthy pregnancies which has been challenged by the development of DNA sequence-based techniques, although it is still open to controversy. Nonetheless, little is known whether different parts of fetal appurtenances contain district microbiome profiles. Here, DNA 16S rRNA sequencing was performed of the amniotic fluid cells (AC), amnion membrane (AM), the placenta of fetal surface (remove the amniotic membrane, PL), maternal blood (MB), and umbilical cord blood (UCB) at V3-V4 hypervariable region from participants with cesarean delivery. Then sequence raw data were followed by taxonomic classification at 97% similarity and diversity analysis at the genus level. The differences and associations among the five tissues were analyzed. At the phylum composition level, the most abundant microorganisms were Proteobacteria in all five tissues, and followed by Firmicutes in AC, AM, and MB groups, Actinobacteria in UCB and Bacteroidetes in PL, respectively. As the maternal-fetal barrier, PL and AM had the lower OUT number and weaker co-occurrence network compared with the other three tissues. At the beta diversity clustering level, the microbiota constituents in the MB and UCB were highly similar; the microbiota profiles of PL and AM were also remarkably alike; AC was immensely different from those two clusters. Therefore, the five tissues were distinctly separated into three clusters. Our study reveals that different pregnancy-related anatomical sites harbor unique microbial compositions and show different degrees of correlation with other tissues.

## 1. Introduction

The human microbiome is an enormous community of microorganisms occupying different body sites of human beings, such as skin, nose, mouth, lung, intestinal tract, and vagina (1–8). ~80% microbes are colonized in the human intestine, playing important roles in nutrient metabolism, immunomodulation, anti-pathogens, free radical scavenging and gut mucosal barrier structure integrity maintenance of their human hosts (9–11). Studies of the Human Microbiome Project have indicated that different human body sites harbor site-specific microbiota. For the reproductive system, the uterus and placenta were traditionally thought to be sterile and microbial invasion of this organ had been associated with adverse pregnancy outcomes. This “sterile womb” paradigm has recently been challenged by new molecular techniques, mainly metagenomics and 16S rRNA gene amplicon sequencing. Several studies have shown that the placenta harbors a unique microbiome, and the microbiomes are altered with different maternal pregnant conditions. Studies of Xinhua Xiao team h that gestational diabetes mellitus (GDM) were associated with placental microbiota alternation. In the placenta, Proteobacteria were increased, and Bacteroidetes and Firmicutes were decreased in women with GDM (12). Their team also found the placental microbiota profile in fetal macrosomia was distinguished from normal infant weight (13), and so did the low birth weight group (14). Moreover, placental microbiota was elucidated to be involved in preterm birth (15, 16) and pre-eclampsia (17). However, it is still controversial about the existence of a universal placental or fetal microbiota, as some researchers showed there was almost a negative culture for bacterial growth from those tissue samples of normal pregnancy. They argued that the 16S ribosomal RNA gene sequencing data might be all related either to the acquisition of bacteria during labor and delivery, or to contamination of laboratory reagents (18–21). However, there are more and more recognitions that ‘non-cultivability’ does not mean “not exist” because there are some challenges to culture bacteria of low abundance in vitro. In healthy term pregnancy, it is also inconclusive whether the amniotic fluid harbors bacteria (10, 22, 23).

Regardless of the controversy, multiple studies showed that the microbiome might play a role in the maintenance of a healthy pregnancy (24, 25). Throughout pregnancy, the microbiome in different body sites undergoes changes associated with metabolic alterations and immunological adaptations (26). The microbiome in district body sites might affect pregnancy outcomes specifically related to its residing niche. Maternal gut microbiota is one of the important factors in the developmental origins of health and disease (DOHaD) concept. Kuang, et al compared the gut microbial composition of gestational diabetes mellitus (GDM) patients and healthy pregnant women by sequencing their fecal samples collected during the second pregnant trimester and found that *Parabacteroides distasonis* and *Klebsiella variicola* were enriched in GDM patients, while *Methanobrevibacter smithii, Alistipes spp., Bifdobacterium spp*., and *Eubacterium spp*. were enriched in normal pregnant women. The results indicated an association between the gut microbiome and GDM status (27). Maternal gut microbial diversity affected the male newborns’ weight and Streptococcus negatively regulated the female newborn’s body height, suggesting the maternal gut microbiota might have sex-specific effects on fetal growth (28). As mentioned above, placental microbiota has been shown a significant association with gestational duration, pregnancy complications, pregnancy outcomes, and infant postnatal development (13, 14, 17, 29–31). A recent study found maternal blood microbiome was also associated with the pregnancy process that Firmicutes and Bacteroidetes were more abundant in maternal blood with preterm birth while Proteobacteria was less prevalent (32). While similar to the placenta, whether there is a live bacterial community in the blood is debatable. Traditionally, blood in healthy humans is thought as a ‘sterile’ environment, and culturing the relevant microbes has rarely been successful. However, the existence of a novel bacteriological system was noted from blood samples taken from healthy humans (33, 34) and was not due to contamination using appropriate and careful controls. Moreover, the previous studies showed the flora in umbilical cord blood were identified as the genus *Enterococcus, Streptococcus, Staphylococcus* belonging to Firmicutes phylum, and *Propionibacterium* belonging to Actinobacteria phylum (35). Another study revealed that blood fractions contain bacterial DNA mostly from the Proteobacteria phylum (more than 80%) but also from Actinobacteria, Firmicutes, and Bacteroidetes, and there are striking differences between the bacterial profiles of the different blood fractions at deeper taxonomic levels (36). All these studies indicate that a diversified microbiome might exist in healthy blood.

Bacteria or their metabolites from the maternal environment might be translocated to the fetus via the bloodstream, and microbes in maternal different body sites might have impacts on the fetus. Therefore, we intend to investigate whether there is any correlation between the microbiome in maternal blood and fetal blood. In addition, we further aim to investigate the profiles and correlations of microbiome among diverse tissues of mother and fetus.

## Material and methods

### Ethics statement

This study was performed with the informed consent of the participants. The experimental design and protocols used in this study were approved by the Third Affiliated Hospital of Guangzhou Medical University Research Ethics Committee (reference ECM 20/02/2019, No.042). The participants in this study were recruited with an informed consent form (ICF) by the Third Affiliated Hospital of Guangzhou Medical University.

In this study, two cohorts of total 28 patients were involved. In cohort 1, the raw data of three volunteers were excluded because the participants had autoimmune diseases or amniotic choritis. Finally, data from 8 participants with normal fetal weight were used to explore the microbiota correlation among diverse tissues. Participants in cohort 2 were all without autoimmune diseases or confirmed infections of the reproductive system, so 17 Participants’ data were analyzed. All the samples including amniotic fluid cells (AC), amnion membrane (AM), the placenta of fetal surface (remove the amniotic membrane, PL), maternal blood (MB, peripheral blood), and umbilical cord blood (UCB) were collected according to SOP during C-sections in the sterile operating room by medical workers complying with all relevant ethics of working with human participants. The samples collected at different time were preserved in liquid nitrogen until sequencing.

### DNA extraction and Polymerase Chain Reaction (PCR)

DNA was extracted with Mag-Bind Soil DNA Kit (M5635-02, Omega) and then detected by 0.8% agarose gel. The bacterial 16S rRNA gene V3–V4 hypervariable region was amplified with the specific forward primer 338F 5’-ACTCCTACGGGAGGCAGCA-3’ and the reverse primer 806R 5’-GGACTACHVGGGTWTCTAAT-3’. Sample-specific 7 bp barcodes were incorporated into the primers for multiplex sequencing. Each PCR reaction contained 5 μl Q5 reaction buffer (5×), 5 μl Q5 High-Fidelity GC buffer (5×), 0.25 μl Q5 High-Fidelity DNA Polymerase (5 U/μl), 2 μl (2.5 mM) dNTPs, 1 μl (10 uM) each forward and reverse primer, 2 μl DNA Template and 8.75 μl ddH2O. PCR amplification was performed as follows: 98 °C for 2 min, followed by 25 cycles consisting of denaturation at 98 °C for 15 s, annealing at 55 °C for 30 s, and extension at 72 °C for 30 s, with a final extension of 5 min at 72 °C. PCR amplicons were purified with Agencourt AMPure XP Beads (A63882, Beckman Coulter, Indianapolis, IN) and quantified using the PicoGreen dsDNA Assay Kit (P7589, Invitrogen, Carlsbad, CA, USA). Then amplicons were pooled in equal amounts and were sequenced on the Illumina MiSeq platform using paired-end 2×300 bp MiSeq Reagent V3 Kit according to the standard protocols.

### Bioinformatics and Statistic Analysis

The sequencing raw data were filtered according to the criteria as previously described (37, 38). Sequence length <150 bp, sequences containing ambiguous bases or mononucleotide repeats>8 bp were excluded. Paired-end reads were assembled using FLASH(fast length adjustment of short reads to improve genome assemblies) (39). After denoising and chimera detection, the remaining high-quality sequences were clustered into operational taxonomic units (OTUs) at 97% of sequence identity by UCLUST (40) and then classified taxonomically by BLAST against the Greengenes Database (41).

### Statistical analysis

Sequence data analyses were mainly performed using QIIME and R packages. Alpha diversity indices, Chao1 richness estimator, ACE metric (Abundance-based Coverage Estimator), Shannon diversity index and Simpson index were calculated in QIIME depending on the OUTs taxonomy. Beta diversity was measured by Euclidean distance metrics and Bray–Curtis distance matrices and visualized via principal component analysis (PCA) and principal coordinate analysis (PCoA) based on the genus-level compositional profiles. The statistical differences of microbiota structure among groups were assessed by PERMANOVA (Permutational multivariate analysis of variance, Adonis) using Bray–Curtis distance and ANOSIM (Analysis of similarities) using weighted unifrac distance metrics by R package “vegan”. Hierarchical clustering of the abundant genera (OUT abundance> 0.05%) was visualized by heatmap and phyla were shown by stacked bar chart to determine microbiota patterns. PLS-DA (Partial least squares discriminant analysis) reveals the microbiota variation among groups using “PLS-DA” function in R package “mixOmics” at the genus level. To construct the co-occurrence networks of microbiota in different tissues, pairwise inter-genus correlations were calculated according to the genera abundance profiles visualized using R package “igraph”.

For all analyses, p<0.05 was considered statistically significant, and significance levels were indicated as follows: ***, p < 0.001; **, p < 0.01; *, p < 0.05.

## Results

### Participant Characteristics

We studied two cohorts of participants. We collected all samples from mothers with cesarean sections in the Third Affiliated Hospital of Guangzhou Medical University. There were 11 participants in the first cohort and 17 in the second cohort. In cohort 1, we chose 8 participants who had no autoimmune diseases or amniotic choritis. In addition, all the infant weights were > 2.5 kg. The mother-child pairs condition details including the maternal pregnancy age, BMI at delivery, gestational duration, and baby weight were provided in Supplementary table 1. For the second cohort, all the participants also had no autoimmune diseases or amniotic choritis, but some parturient women suffered from gestational diabetes or preeclampsia so we mainly showed the results of cohort 1 and put the results of cohort 2 as a repeat to validate the conclusion. Samples from AC, AM, PL, MB, and UCB were sequenced on the Illumina MiSeq platform. An average of ~36,000 reads were then analyzed for each sample.

### Taxonomic composition and alpha diversity of the microbiota from five different tissues

To analyze the taxonomic composition of the microbiota in the five tissues, we aligned the 16S rRNA sequences against the Greengenes Database (41). On average, 35,953 16S rRNA sequence reads were obtained, which are sufficient to detect the microorganisms in the 40 samples as shown by the rarefaction curve (Fig. S1). Proteobacteria were the most abundant phylum among all the tissue samples with an average of 85.355% (Fig. 1A). Then followed by Actinobacteria, Firmicutes, Bacteroidetes, and Thermi which were all with an abundance greater than 1% on average (Fig. 1A). At the phylum level, we concluded that the MB and UCB had the similar microbial compositions, for example, the levels of Firmicutes (P=0.46) and Bacteroidetes (P=0.78) were almost identical in these two tissues. AM and PL had very similar compositions, such as the almost same levels of Actinobacteria (P=0.95), and similar levels of Bacteroidetes (P=0.25) (Fig. 1A). At the genus level, Cupriavidus and Burkholderia were the two most abundant bacterial genera belonging to Proteobacteria among all the samples (Fig. 1B). As shown by the genus heatmap, the AM and PL had highly similar composition pattern, and so did the MB and UCB. AM is the most inner layer of the placenta making up the maternal-fetal barrier, close to PL (placenta, remove the amniotic membrane), therefore we hypothesized that the high similarities of microbiome between AM and PL might be because of the adjacent physiological location and function. Interestingly, the microbiome profiles in MB were very similar to that of UCB. Thus, each body site harbors unique microbiomes although there were little variations between different individuals.

**Figure 1.**
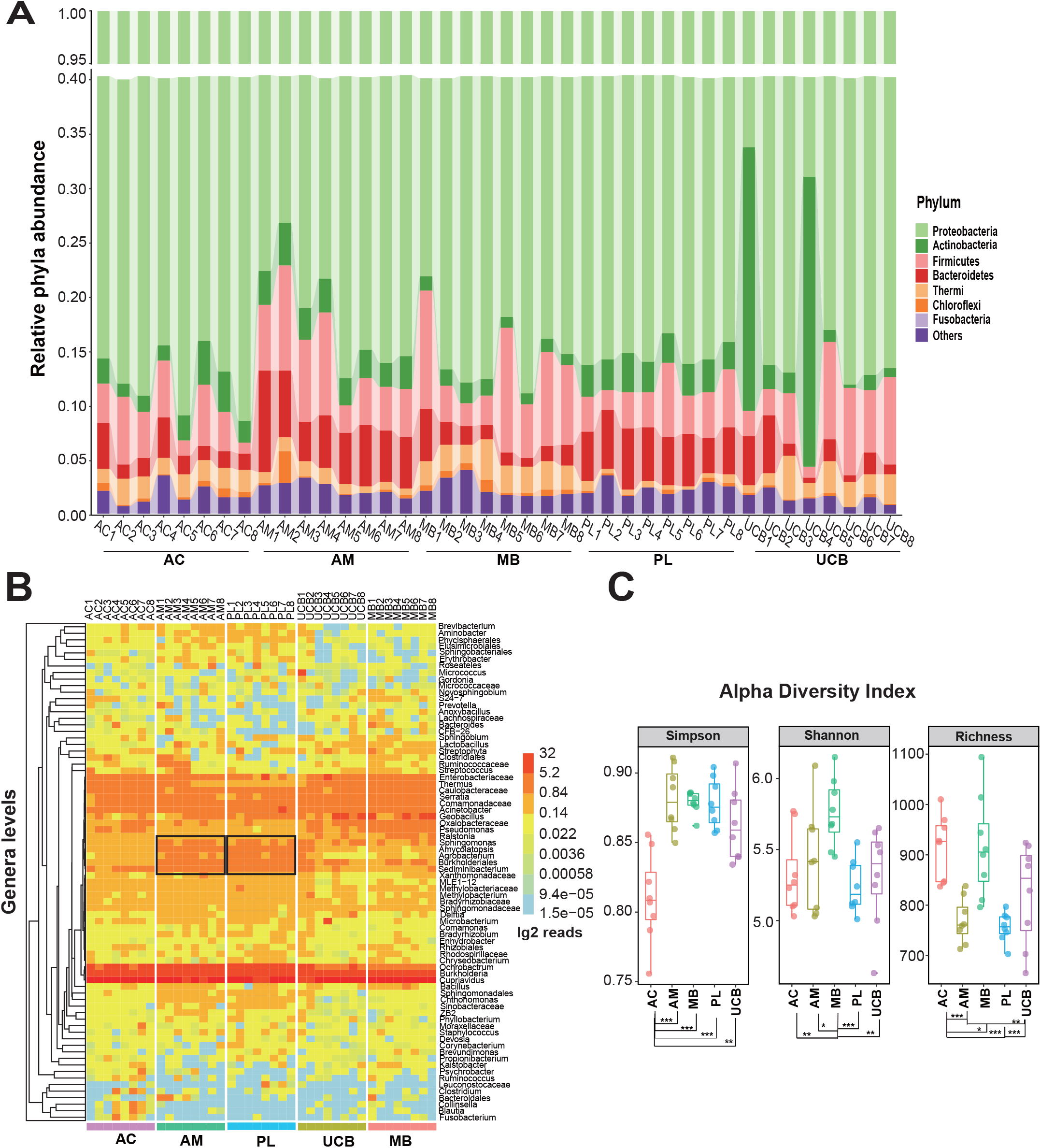
Taxonomic composition and richness of the five tissues microbiome. **(A)** Seven phyla were identified with an average relative abundance greater than 0.35% among all samples. **(B)** Heatmap based on top 76 genera among five tissues. The reads number is indicated by a color gradient from light blue (low) to red (high). **(C)** Alpha diversity was shown as Simpson whose value is negatively correlated with α-diversity and Shannon whose value is positively correlated with α-diversity. Richness was indicated as OUT number.

Besides the taxonomic composition diversity, the microbiota evenness and richness also varied among different tissues. In accordance with the maternal-fetal barrier functions, PL and AM had the lower OUT number compared with the other three tissues (Fig. 1C).

### Differences in microbial community compositions among five groups

The taxonomic composition histogram (Fig.1A) showed the phyla percentages with abundance greater than 0.5%. The relative abundances of four phyla including Thermi, Acidobacteria, Bacteroidetes, and Chloroflexi were significantly different among the five groups (Fig. 2A). The differences in genus level were performed by STAMP software between every two groups. The microbes between PL and AM, UCB and MB were highly similar as shown in the genus heatmap of Figure 1B. Then, we further compared the AC with PL, AC with MB, and PL with MB (Fig. 2B). Genus differences among other tissues (UCB and MB, UCB and AM, UCB and AC, UCB and PL, AM and MB, AM and PL, AM and AC) were shown in Supplementary figure 2 (Fig. S2). Differential genera in AC, MB, and PL were *Cupriavidus, Enterobacteriaceae, Serratia, Burkholderia, Ochrobactrum, Comamonadaceae, Burkholderiales, Oxalobacteraceae, Pseudomonas, and Agrobacterium* which are all belonging to Proteobacteria phylum; *Geobacillus* belonging to Firmicutes; *Thermus* belonging to Thermi and *Sediminibacterium* belonging to Bacteroidetes (Fig. 2B), the genera differences were consistent with the differences on phyla level (Fig. 2A).

**Figure 2.**
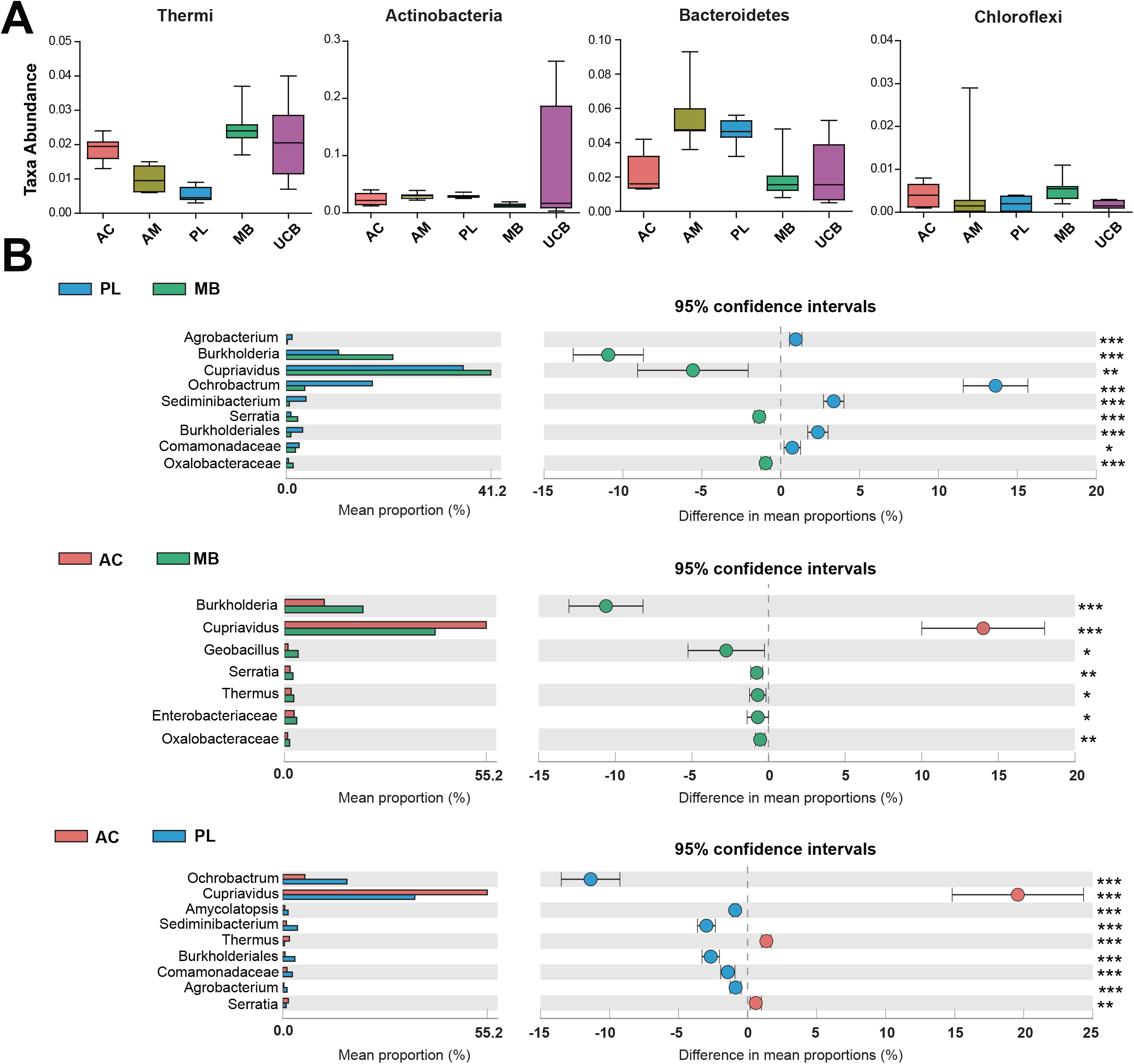
The microbe differences among tissues at phylum and genus levels. **(A)** Four phyla with an average relative abundance greater than 0.1% were identified that their relative abundances were different among the five groups. **(B)** Difference analysis of genus levels between two groups in AC, MB, and PL tissues with two-sided Welch’s t-test on STAMP platform. Genera with significant differences and an average relative abundance greater than 0.5% were shown.

Then we re-organized the PL and AM to one group (named Placenta), and the UCB and MB as another group (named Blood) to deeply explore the significance of changes in bacterial communities among the three related tissues AC, Placenta, and Blood by LEfSe. Several discriminative taxa were identified with high proportions in Betaproteobacteria and Alphaproteobacteria classes between AC and Placenta groups (Fig. 3A~3B). While, Placenta and Blood displayed no significant divergences by LEfSe, neither did AC and Blood groups (data were not shown). These results indicated that the microbial community compositions between AC and Placenta were more separate.

**Figure 3.**
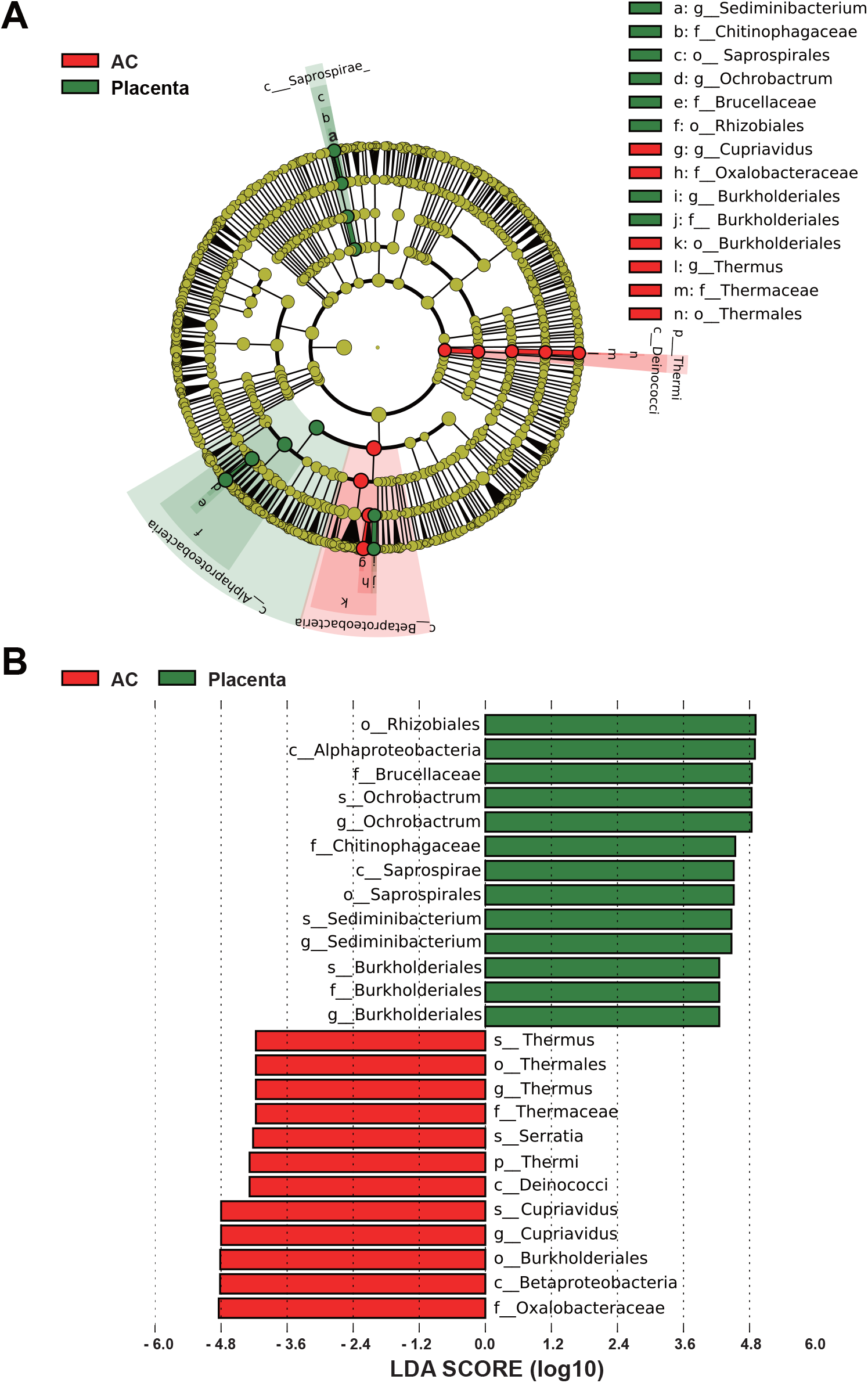
The microbe differences from LEfSe analysis between AC and Placenta. **(A)** PL and AM were recognized as one group (Placenta). Taxa (Green) enriched in the Placenta group and taxa (red) in AC tissue. **(B)** Placenta-enriched taxa were shown with a positive LDA score (green) and AC-enriched taxa harbored a negative score (red). Only the taxa meeting the condition of a logarithmic LDA score significant threshold>2, P < 0.05 were presented.

### Microbiota structure differences and associations in five different groups

Our previous data showed that the microbiota compositions varied among different tissues (Fig. 1A~1B), we further performed the beta diversity clustering analysis with PCA and PCoA by PERMANOVA test. Beta diversity measures the between-group differences and relevancies. PCA is calculated depending on the Euclidean distance matrices and PCoA is based on the Bray–Curtis distance matrices. There was a notable separation among five groups sampled at each body site depending on the PCoA results (R=0.414, P=0.001) (Fig. 4A). Each group separated from other groups based on PC2 direction, while in PC1 axial direction, UCB and MB, PL and AM had the much-closed distribution (Fig. 4A) in accordance with the Bacterial community results (Fig. 1A~1B). Moreover, the width of the link line between two group center nodes represents the degree of correlation indicating that AM and PL, UCB and MB were highly related, respectively. Interestingly, microbiota structure from AC was more related to that of UCB compared with that of MB, maybe it is because AC and UCB all come from fetus tissues (Fig. 4A). Consistent with the LEfSe results (Fig. 3A~3B), AC almost had no correlation with AM/PL (Placenta). The PCA vision also showed similar results (Fig.4B). Importantly, the PCoA results from cohort 2 showed similar relations among five tissues (Fig.S3), and microbiome in MB and UCB had no significant differences (R=0.003, P=0.98). ANOSIM dissimilarity comparisons between every two groups further corroborated the PCoA conclusion that AC microbiota profiles were obviously different from that of the other four groups, and PL/AM were distinguished with MB/UCB (Fig. 4C), but no significant differences occurred between PL and AM, or UCB and MB, respectively (Fig.S4).

**Figure 4.**
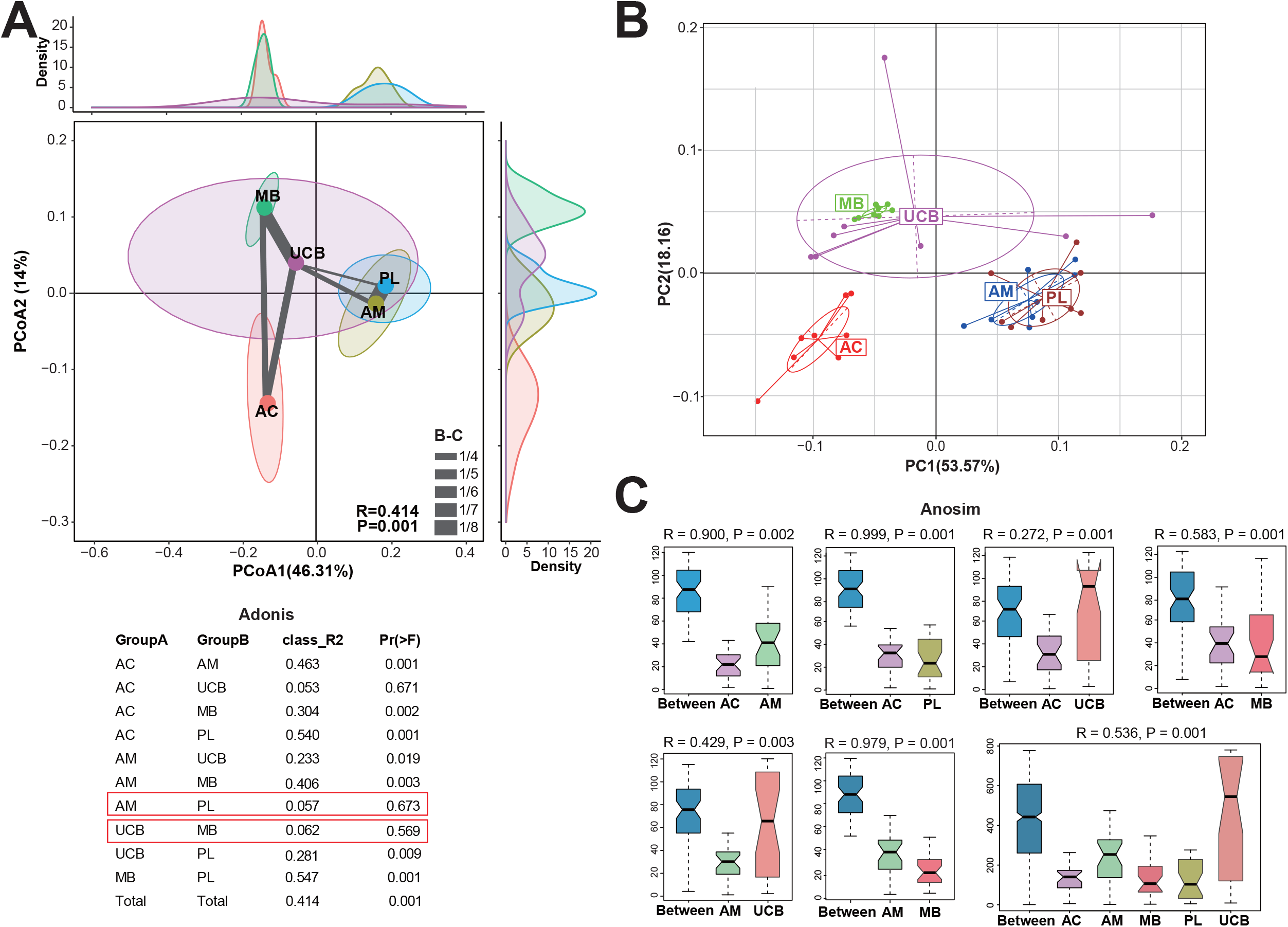
Beta diversity of the microbiota in five tissues. **(A)** Principal coordinate analysis (PCoA) based on Bray–Curtis distance matrices according to the genus-level compositional profiles in five tissues. The correlation values in every two groups were indicated as the Bray-Curtis distance matrices. Significant differences in microbiota structure among groups were assessed by Adonis also based on Bray-Curtis distance matrices. **(B)** Principal component analysis (PCA) was performed with Euclidean distance metrices among five tissues on genus levels. **(C)** Analysis of similarities (ANOSIM) between two low-correlative groups and five groups was performed based on the weighted-unifrac distance metrics of OUT profiles.

Furthermore, we used a redundancy analysis (RDA) plot to explore complex associations between community composition and various explanatory variables, the results were consistent with PCA/PCoA analysis (Fig. 5A). Conjoint analysis of RDA1 (37.82%) and PC1 (19.57%) principal components showed that the AM and PL microbiota structures were highly associated (Fig. 5B). The width of link line in Figure 4A showed that the microbiome of AC was more relevant to that of blood tissues (MB and UCB), and the results of figure 5A also showed the microbiota of AC was more closed to that of MB.

**Figure 5.**
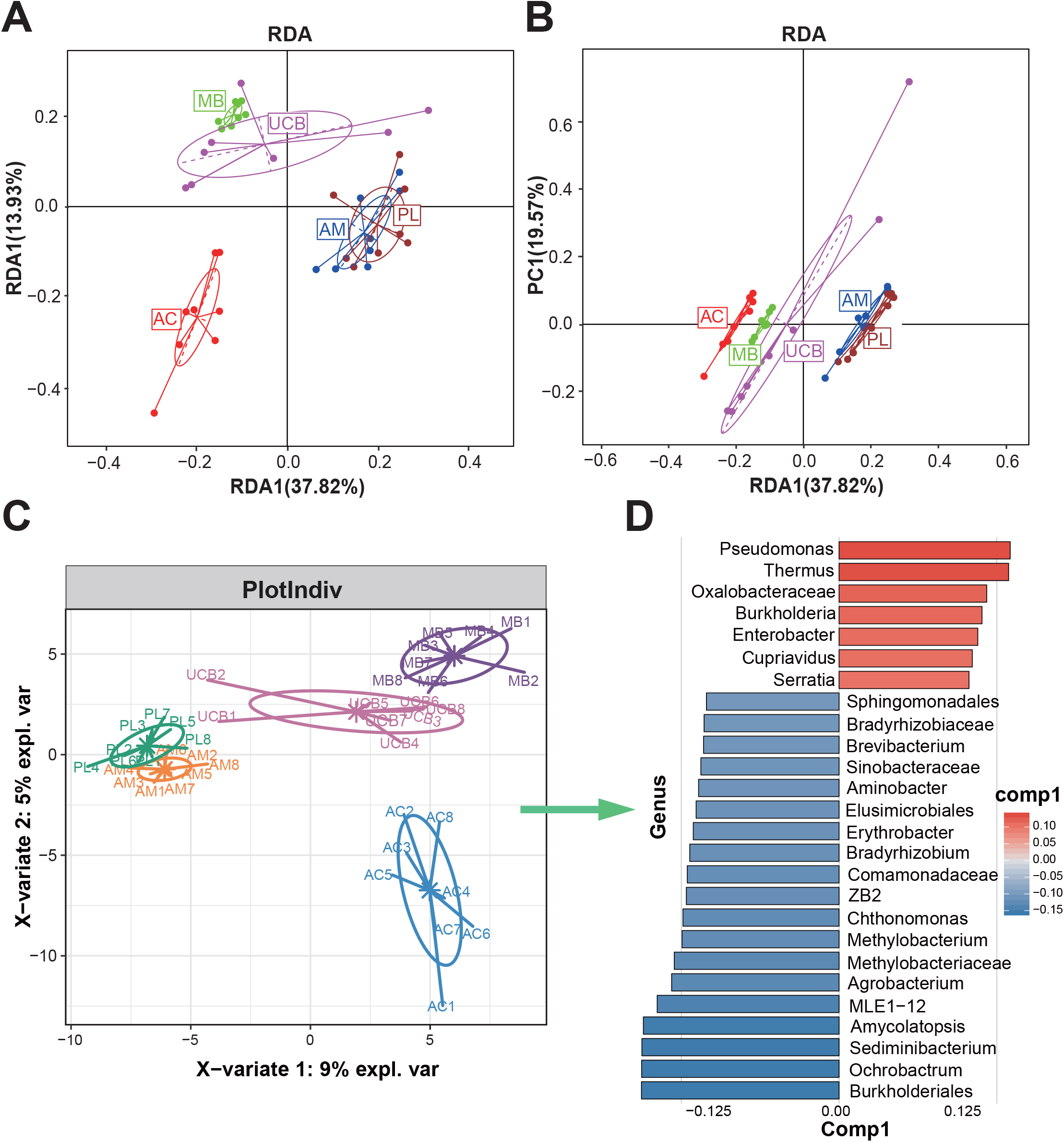
Samples separate of different body parts based on genus composition profiles. **(A)** Redundancy analysis (RDA) of genera with average relative abundance greater than 0.1% among five groups showed separation of samples by body sites. **(B)** RDA1 and PC1 conjoint analysis was performed to separate groups on genus levels. **(C, D)** The Sparse Partial Least Squared–Discriminative Analysis plot illustrated a clear separation in five tissues based on the genera of greater than 0.1% relative abundance. The related contribution plot illustrated taxa associated with the fetal-maternal interface tissues.

Next, we identified the most discriminative taxa, which can best characterize microbial compositions of five tissues. Sparse partial least squared–discriminative analysis (sPLS-DA) was conducted on the abundant genera average greater than 0.1% proportion. *Pseudomonas, Thermus, Oxalobacteraceae, Burkholderia, Enterobacter, Cupriavidus*, and *Serratia* were found to best characterize the microbial genera compositions in the blood (MB and UCB) and Amniotic fluids (AC). While, *Sphingomonadales, Bradyrhizobiaceae, Brevibacterium, Sinobacteraceae, Aminobacter, Burkholderiales, Ochrobactrum, Sediminibacterium, Amycolatopsis, MLE1–12, Agrobacterium, Methylobacteriaceae, Methylobacterium, Chthonomonas, Bradyrhizobium, Erythrobacter, Elusimicrobiales, ZB2*, and *Comamonadaceaeat* were the characterized genera at placental tissues (PL and AM) (Fig. 5C~5D).

### Microbial co-occurrence network analysis

Since microbiota varies from person to person, so it is important to investigate the coordinated interactions of microorganisms colonized in the same body sites among the 8 participants. We constructed co-occurrence networks of the microorganisms by calculating the pairwise inter-genus correlations based on genera abundance profiles of 8 samples in every group. We found that the strength of the microbial co-occurrence was significantly greater in AC or UCB groups which are originally from fetus tissues, suggesting that the microorganisms from fetus tissues are very steady and coordinated (Fig. 6 and Fig. S5). This might be because the fetus is in a relatively stable microenvironment during pregnancy. Conversely, bacterial profiles in MB had the lowest co-occurrence, which may reflect the various physical conditions of adults.

**Figure 6.**
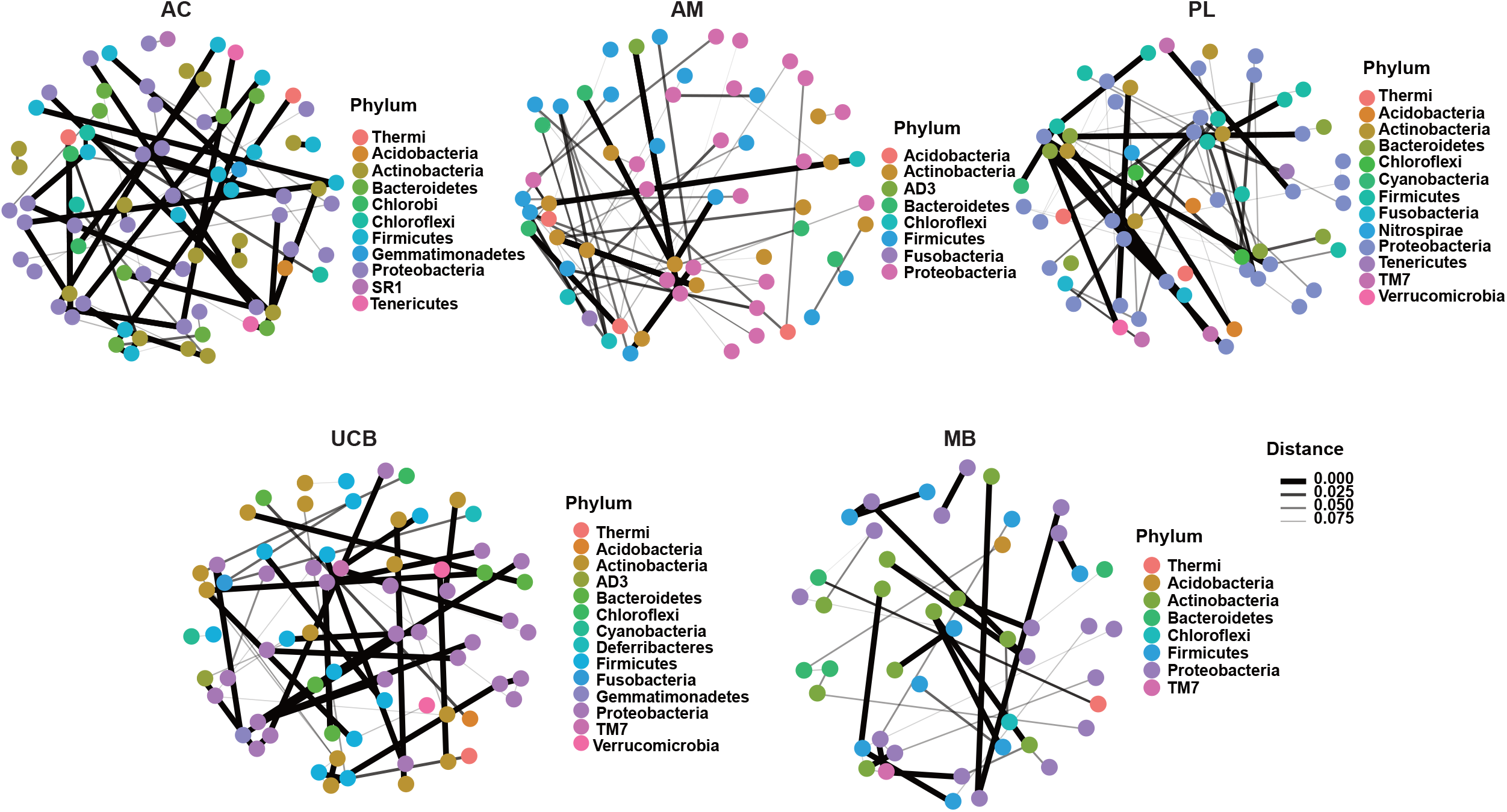
Microbial co-occurrence network analysis. Co-occurrence networks were constructed using Bray–Curtis distance matrices less than 0.1 based on the genera relative abundance profiles among the five groups. The smaller the distance, the stronger the co-occurrence correlation. Each node represented a genus and genera belonging to the same phylum were shown in one color.

## Discussion

The placenta plays important roles in sustaining fetus survival as both a lifeline and a guardian, it shuttles oxygen, nutrients, and immune molecules from the mother to her fetus. Placenta also serves as a barrier against infections. For a long time, the placenta and even the womb were thought to be sterile unless something went wrong during pregnancy. However, more and more studies have suggested the existence of the placental microbiome, which might even be a crucial part of pregnancy, could have an important role in shaping the developing immune system (42). Therefore, it is worth exploring the microbiota profiles in different tissues at the maternal-fetal interface.

In this study, we performed bacterial 16s rRNA sequencing from several reproduction-related tissues. Clear and distinct microbiomes were identified in every tissue. Among those tissues, the microbiomes in MB and UCB were highly similar (P=0.569), and were separate from that in the placenta although the placenta is infiltrated by blood. Meanwhile, PL and AM harbored highly alike microbiomes. MB and UCB have functional similarity, PL and AM are anatomically and functionally related. Therefore, our studies showed that different tissues in the maternal-fetal interface harbor distinct microbiomes, and the profiles of the microbiomes are related to their anatomical position or function.

Previous research has shown that maternal microbiota in other body sites such as oral (43–45)and gut (26, 27) could affect the pregnancy processes and outcomes. The oral flora can be capable of oral-uterine transmission during pregnancy confirming the transferability of colonized flora. Studies have also suggested that the maternal microbiome during pregnancy might have a key role in preventing an allergy-prone immune phenotype (46) or influencing neonatal immunity (31) of the offspring. In addition, the maternal microbiota might have a role in mother-infant interaction and perinatal depression (47). However, the mechanisms remain unclear. Microbiota transfer from mother to fetus would mediate the maternal impact on infants even till childhood. So far, studies about the microbiomes in maternal and umbilical cord blood are scarce. Our finding of the highly similar microbiome profiles in MB and UCB suggests that the microbiota in MB or UCB may be related to blood functions. The data from 17 participants of cohort 2 also showed that the microbiome profiles between MB and UCB were highly similar, demonstrating the strong relevance of microbiome in mother and fetus. This strong correlation suggests that the microbiome might be a possible mediator for mother-to-infant epigenetic heredity.

Compared with the other three tissues, PL and Am have lower OUT numbers and weaker co-occurrence networks, coinciding with their role as barriers. In terms of taxonomic composition, our results were slightly different from another research with placenta samples collected from Beijing, China. They found that Proteobacteria was the most abundant phyla and then followed by Firmicutes in microbiota from PL and AM (13). Since our samples were collected from the southern part of China, the minor differences in microbiota composition might be related to the climate and diet dissimilarity between Guangdong province and Beijing. How do the climates and diets affect the composition of the microbiome in the placenta? Studies have shown that oral dysbiosis is related to adverse pregnancy outcomes, suggesting there might be crosstalk of microbiota between the placenta, oral, and intestine. However, little is known about the microbiota mobility and exchange between mother and fetus.

The strengths of our study include its system and two cohorts design, with paired mother-baby tissues and fetal appurtenances, which allowed us to investigate the microbiota profiles in various body sites. However, the sample size in this study was limited, we will recruit more participants for large-scale studies to investigate the relationship between maternal microbiota and offspring development. Another limitation was the lack the detection of bacteria commonly found in the environment. In fact, we had run a “kit contaminant” control using ddH2O as the amplification template while the library construction was unsuccessful. Still, the microbiome profiles exhibited significant differences among different tissues, which cannot be solely due to contamination.

Collectively, our data showed that different tissues in the maternal-fetal interface harbor clear and distinct microbiomes. Our data support that the fetus harbors unique microbiomes in the blood and shed skin cells before birth. The microbial co-occurrence is significantly greater in AC and UCB tissues which are originally from the fetus indicating that the fetal microorganisms might be more steady than in adult mothers. We speculate it might be because the fetus is in a relatively protected microenvironment during pregnancy. It sounds paradoxical but interestingly that the fetus’s microbiome is affected by maternal flora while resisting maternal variability. Probably, this is an important mechanism of healthy pregnancy sustaining. Therefore, our data systematically reveal the correlations of microbiota among different reproductive tissues and observe a possible role of microbiota in mother-to-baby crosstalk. In addition, our study opens up opportunities, whereby maternal microbiota interventions may be beneficial for infant health care after birth through modulating their microbiota when in the maternal uterus.

## Availability of data and materials

The sequences generated and analyzed during the current study were uploaded to the NCBI Sequence Read Archive (SRA) data depository, with the project number PRJNA635545.

## Author contributions

Xiaopeng Li, Bolan Yu, and Min Fang conceived and designed the experiments and wrote the draft. Xiaopeng Li analyzed the data and Wei Jiang contributed some suggestions. Guihong Liu and Lijuan Dai collected the samples. All authors approved the submission of this version.

