## Supplemental figures and extended methods for "Different tissues in the maternal-fetal interface harbor distinct microbiomes showing associations related to their anatomical position or function"

### Supplementary table 1. Maternal and newborn characteristics.

| Volunteers ID | Pregnant age (year) | BMI at delivery | Gestational duration (week + day) | Infants weight (kg) |
|---|---|---|---|---|
| 1-1 | 47 | 25.7 | 33+6 | 2.72 |
| 1-2 | 22 | - | 36+6 | 2.58 |
| 1-3 | 27 | 18.73 | 40 | 2.85 |
| 1-4 | 37 | 27.05 | 38+5 | 2.50 |
| 1-5 | 31 | 20.43 | 35+2 | 2.62 |
| 1-6 | 30 | 25.2 | 38+5 | 2.99 |
| 1-7 | 25 | 21.09 | 33+1 | 2.52 |
| 1-8 | 40 | 23.7 | 39+2 | 3.63 |

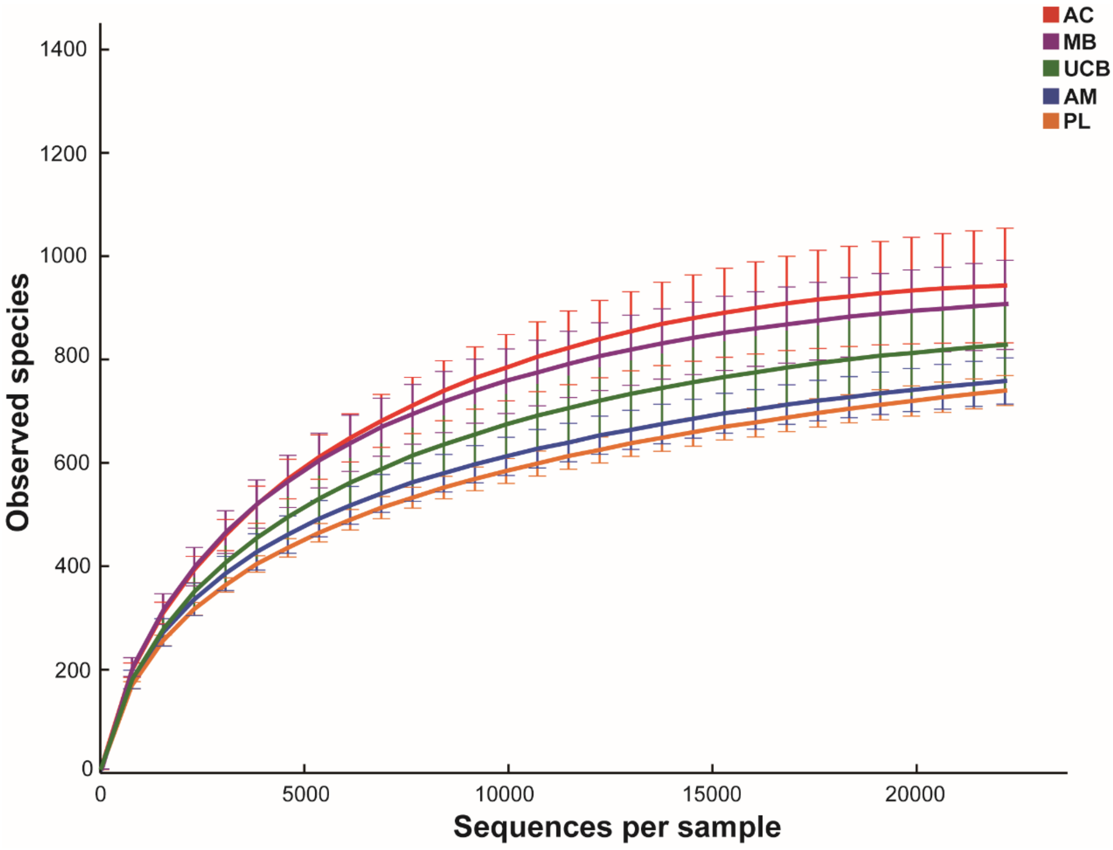


**Supplementary Figure 1.** Rarefaction curve of 16S rRNA sequences in five groups. Data are represented as mean ± sem.

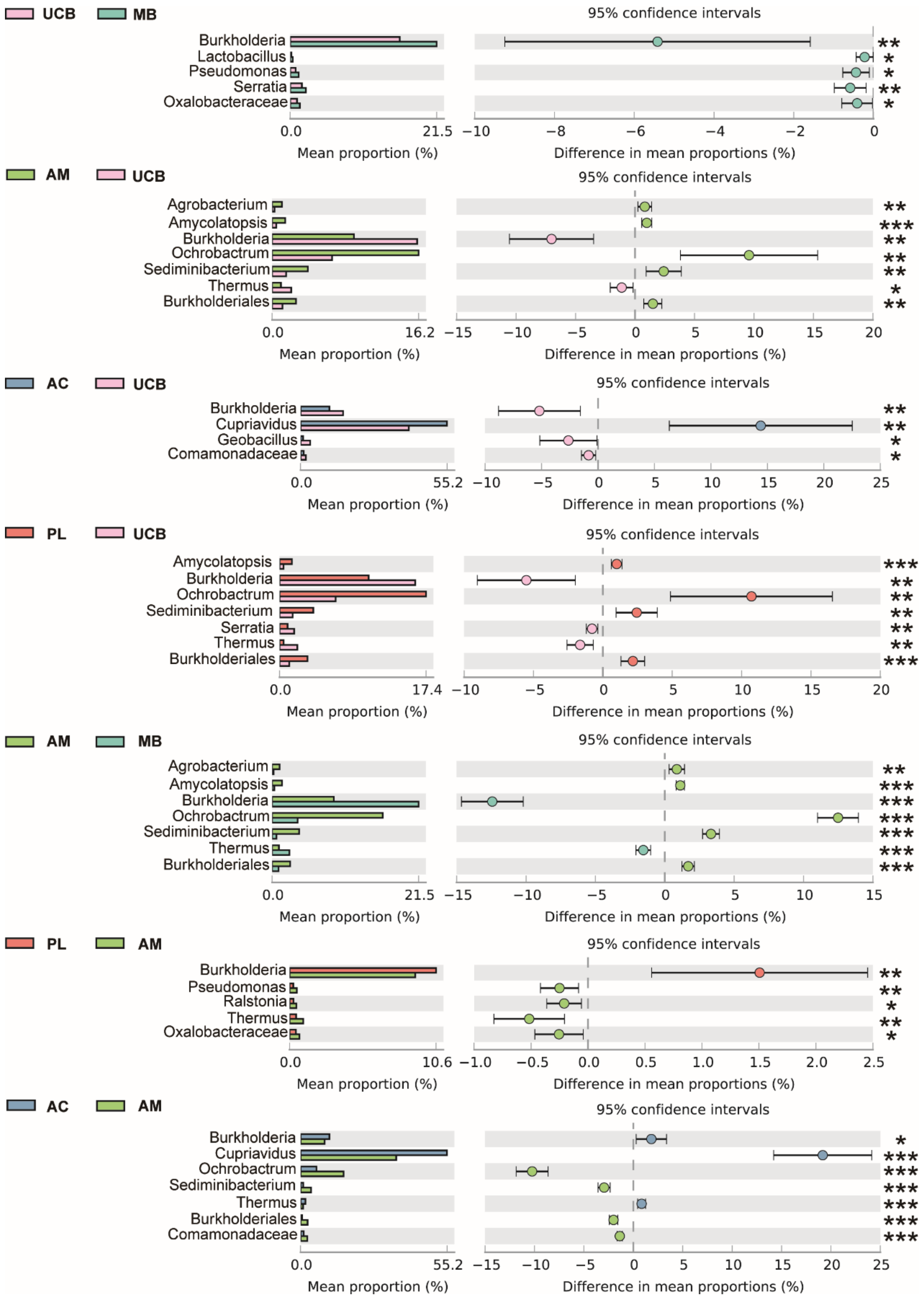


**Supplementary Figure 2.** Difference analysis of genus levels between two groups in five tissues with two-sided Welch's t-test on STAMP platform. Genera with significant differences and an average relative abundance greater than 0.5% were shown.

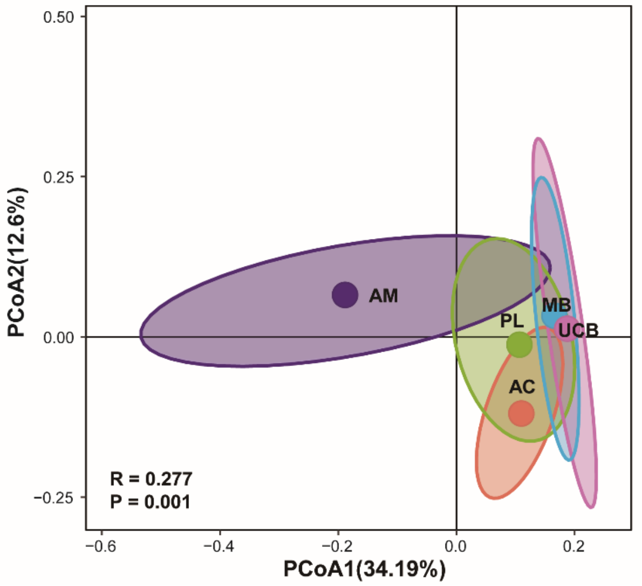


**Adonis**

| GroupA | GroupB | class_R2 | Pr(>F) |
|---|---|---|---|
| AC | AM | 0.124 | 0.001 |
| AC | UCB | 0.108 | 0.004 |
| AC | MB | 0.073 | 0.017 |
| AC | PL | 0.020 | 0.779 |
| AM | UCB | 0.142 | 0.001 |
| AM | MB | 0.125 | 0.001 |
| AM | PL | 0.048 | 0.17 |
| UCB | MB | 0.003 | 0.98 |
| UCB | PL | 0.052 | 0.093 |
| MB | PL | 0.021 | 0.8 |
| Total | Total | 0.277 | 0.001 |

**Supplementary Figure 3.** Principal coordinate analysis (PCoA) based on Bray–Curtis distance matrices according to the genus-level compositional profiles in five tissues of other 17 volunteers. Significant differences of microbiota structure among groups was assessed by Adonis also based on Bray–Curtis distance matrices. **(B)** Analysis of similarities (ANOSIM) between two groups was performed based on the weighted unifrac distance metrics of OUT profiles.

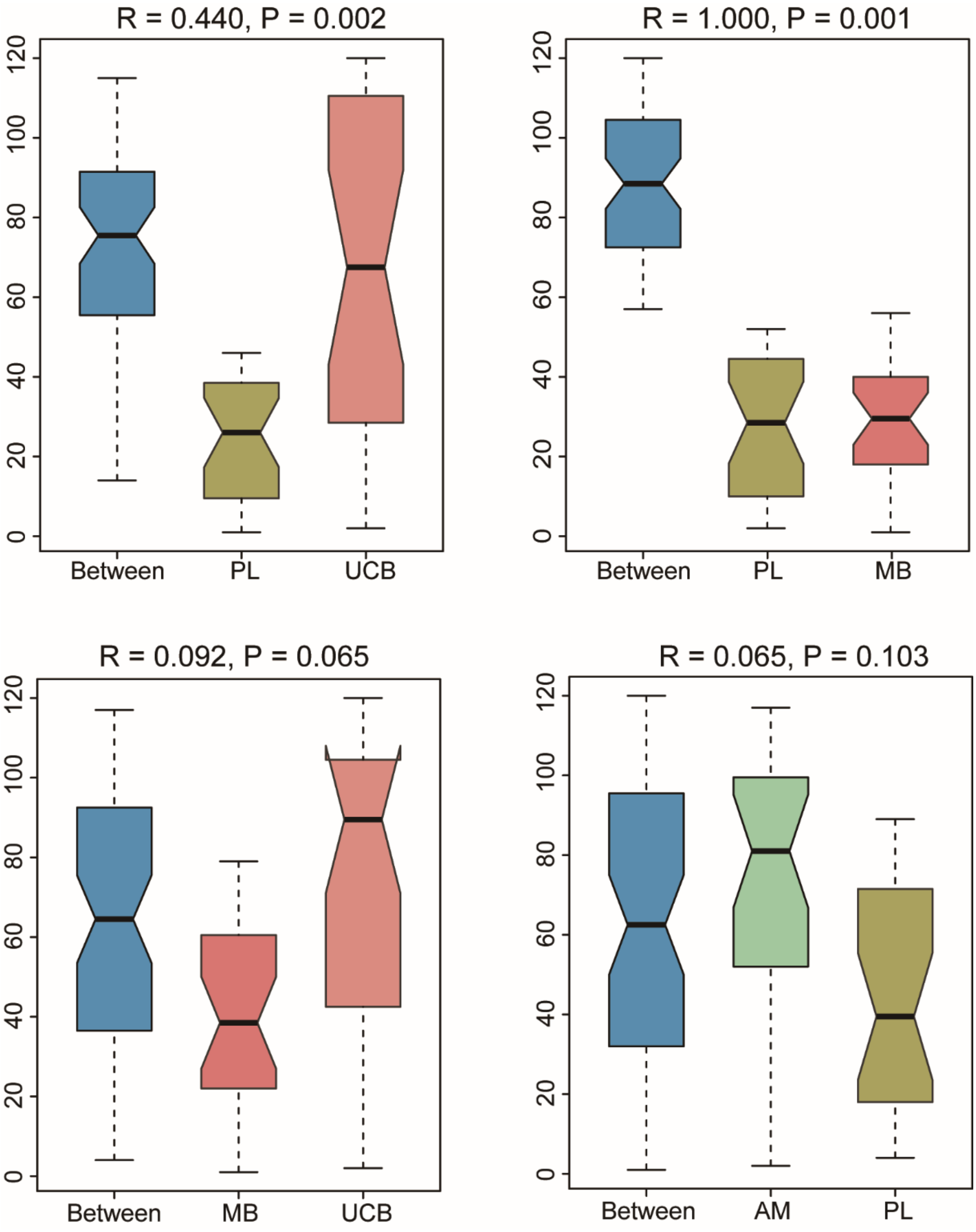


**Supplementary Figure 4.** Analysis of similarities (ANOSIM) between two groups was performed based on the weighted unifrac distance metrics of OUT profiles.

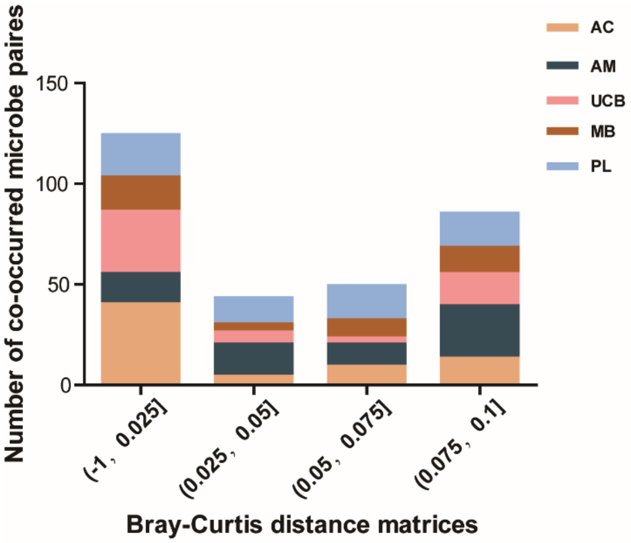


**FDR test**

| GroupA | GroupB | P value |
|---|---|---|
| AC | AM | 0.0003 |
| AC | UCB | 0.2878 |
| AC | MB | 0.2725 |
| AC | PL | 0.0083 |
| AM | UCB | 0.0054 |
| AM | MB | 0.1798 |
| AM | PL | 0.2758 |
| UCB | MB | 0.1798 |
| UCB | PL | 0.0083 |
| MB | PL | 0.4516 |

**Supplementary Figure 5.** The strength distribution of microbial cooccurrence of microbiome in five tissues. The strength was determined based on the Bray-Curtis distance matrices less than 0.1 and the significant differences were analyzed by FDR test. The microbial cooccurrence was higher in AC and UCB tissues and there was no significant difference between these two tissues.
